# Pyrethroid insecticides susceptibility of *Stomoxys calcitrans* (Linnaeus) and *Stomoxys indicus* Picard (Diptera: Muscidae) from cattle farms in southern Thailand

**DOI:** 10.1101/2021.12.20.473418

**Authors:** Sokchan Lorn, Krajana Tainchum, Pitunart Nusen, Anchana Sumarnrote, Theeraphap Chareonviriyaphap

## Abstract

The susceptibility to six pyrethroid insecticides (permethrin, deltamethrin, alpha-cypermethrin, cypermethrin, lambda-cyhalothrin and bifenthrin), each at the recommended concentration, was evaluated for the two stable fly species *Stomoxys calcitrans* (Linnaeus) and *Stomoxys indicus* Picard, through tarsal contact using a World Health Organization (WHO) cone bioassay procedure. The field populations of stable flies were collected from three study sites (Songkhla, Phattalung and Satun provinces) in Thailand. The stable flies were exposed to insecticide-treated filter paper for 30 min and their knockdown counts at 30 min and 60 min and mortality counts at 12 hr and 24 hr were recorded. The *S. calcitrans* and *S. indicus* in Songkhla and Phattalung populations were moderately susceptible to pyrethroids for 24-hr mortality. Nonetheless, the Satun population of *S. indicus* was completely susceptible to permethrin with 100% mortality and the lowest susceptible to deltamethrin and bifenthrin. The results indicate a generally low susceptibility of stable flies to pyrethroids in the southern provinces of Thailand.

## Introduction

Stable flies (Diptera: Muscidae) are known as common blood-sucking insect pests worldwide. They are among the most important globally recognized pests that impact livestock and wildlife health There are two common species for diurnal and crepuscular activity abundance in Thailand namely, the cosmopolitan species *Stomoxys calcitrans* (Linnaeus 1758) and *Stomoxys indicus* Picard, respectively (Foil and Hogsette 1994; Tainchum et al. 2010, Changbunjong et al. 2012, Keawrayup et al. 2012, Lorn et al. 2020). Stable flies inflict painful bites that annoy, stress and interrupt feeding to livestock, causing loss of blood, lactation and weight loss (Campbell et al. 1987). In 1994, Foil and Hogsette (1994) stated that a 28% weight loss was recorded in livestock attributed to the direct feeding and various defensive behaviors against the stable fly bites all day long. These insects are biological vectors and mechanical carriers of particular pathogens, which cause problems in animal production systems (Changbunjong et al. 2012, Baldacchino et al. 2013). Moreover, stable fly bites can be a considerable annoyance to humans, especially along the beaches in West Florida where they can severely affect the tourist industry. In the USA, economic losses caused by stable flies were estimated at US$432 million in 1991 (Kunz et al. 1991) and the cost for stable fly control was estimated at over US$100 million (US$9.80 per head) in 1993. The estimated economic losses in the United States, caused by stable flies, were US$1 billion per year (Taylor and Berkebile 2006, Taylor and Berkebile 2008), and the reduction of animal productivity cost over US$2.2 billion in 2012 (Taylor et al. 2012).

Stable flies are controlled by many methods, but the main method relies on insecticides. However, several studies have reported that insecticides did not provide effective stable fly control (Marcon et al. 1997, Pitzer et al. 2010, Reissert-Oppermann et al. 2019). The reduced susceptibility to insecticides in a fly population is explained by a selection effect that leads to resistant fly strain (Cilek and Greene 1994). In 1958, insecticide resistance of stable fly to DDT, dieldrin, dilan and methoxychlor has been recorded (Brown et al. 1971). The resistance in a stable fly to pyrethroid (permethrin) and organophosphate (dichlorvos and stirofos) was first reported in Kansas (Cilek and Greene 1994). A few years later, stable flies from southeastern Nebraska, USA, showed low susceptibility to permethrin, stirofos, methoxychlor (Marcon et al. 1997) and a population from Florida was also reported to have low susceptibility to permethrin (Pitzer et al. 2010). Then *S. calcitrans* isolated in southwestern France was resistant to five pyrethroids (cypermethrin, deltamethrin, fenvalerate, lambda-cyhalothrin and permethrin) (Salem et al. 2012). Recently, Olafson et al. (2019) reported that stable fly from Nakhon Ratchasima province, Thailand, was resistant against pyrethroid.

The pyrethroid insecticides are typically highly toxic to insect pests but harmless to mammalians. Over the past decade, pyrethroids have been used in Thailand to control insect pests in both the agricultural and public health sectors. There is a series of registered products, from pyrethrin to synthetic pyrethroid insecticides, launched for the public as well as residential uses that may induce urban runoff and water contamination, as well as deposits in the soil. However, there is no prior report on the susceptibility of stable flies in Thailand to insecticides, or on the level of resistance in stable fly populations. The current study addresses these gaps in knowledge, especially regarding pyrethroid insecticides that are widely used in and around agricultural farms and by households.

## Materials and Methods

### Stable fly sampling

Adults of two common local species of stable flies in southern Thailand (Lorn et al. 2020), namely *Stomoxys calcitrans* and *Stomoxys indicus,* were collected from three localities in Southern Thailand (Songkhla: SON, Phattalung: PHA and Satun: SAT provinces) during September-November 2018 (Table 1; Fig. 1). These three study sites have been known for organic cattle farm learning centers, and on two farms that belong to universities insecticide treatments for pest control are applied, while one farm pursues organic farming practices. Adults of both sexes of the stable flies were captured between 0800 and 1800 hours, with a sweeping net and a mouth aspirator and they were kept alive and transferred to clean insect cups and later to insect cages. Morphological identification of species was individually performed under a stereomicroscope, following the taxonomic keys (Changbunjong et al. 2012, Baldacchino et al. 2013) for sorting by species to *S. calcitrans* and *S. indicus*. The collected alive specimens were kept in net cages and were provided ad libitum as nutrition — 100% organic honey solution and water-soaked on a cotton ball. The cages were placed in an insectary at 27°C and 80% RH in the Pest Management Laboratory, Faculty of Natural Resources, Prince of Songkla University, Hat Yai Campus, Songkhla Province, Thailand, for a couple of days for their blood meal digestion before bioassay test.

**Fig 1.**
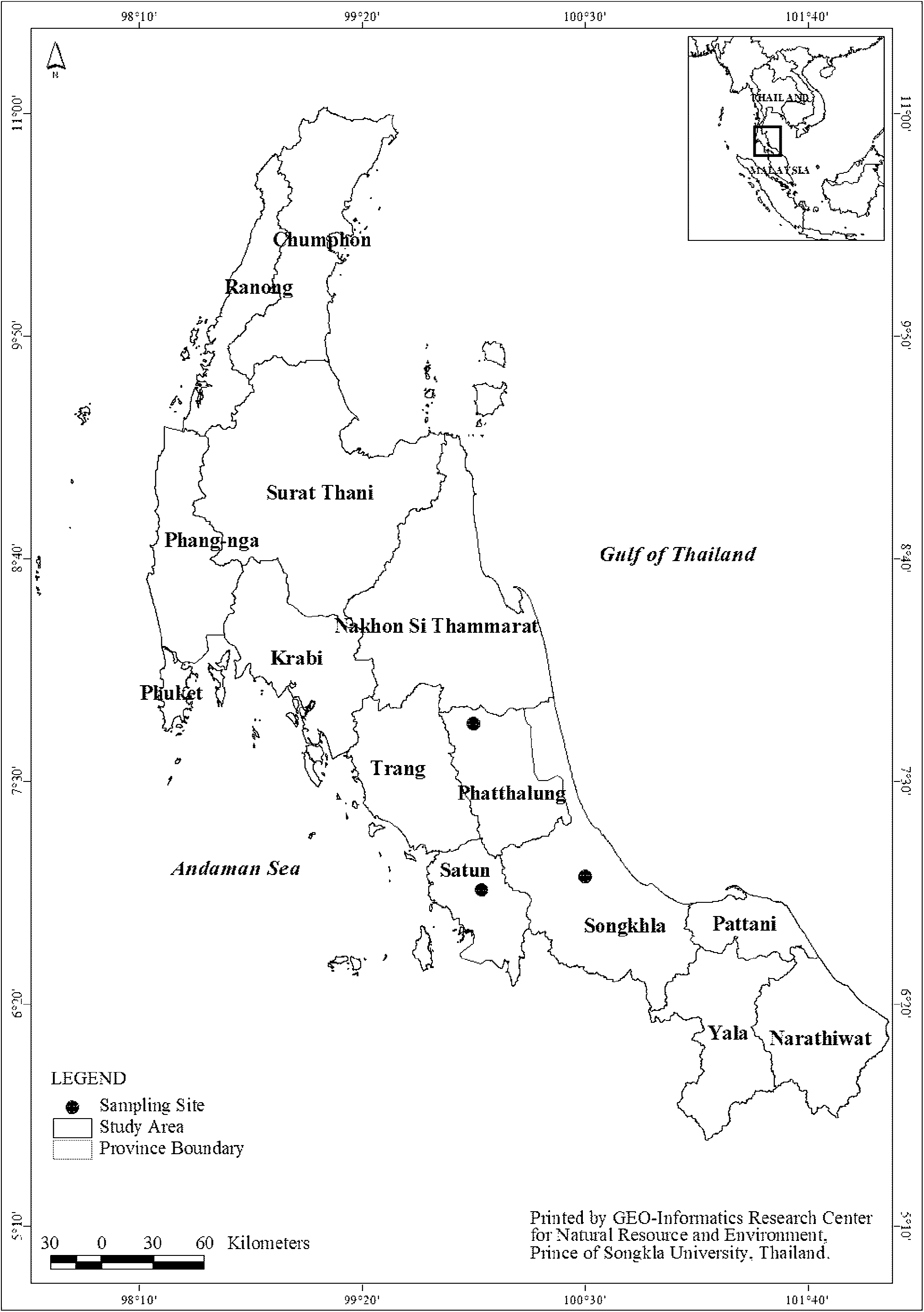
Collection sites of *Stomoxys calcitrans* and *Stomoxys indicus* were located in 3 provinces; Phatthalung, Satun, and Songkhla, Southern Thailand

**Table 1.**
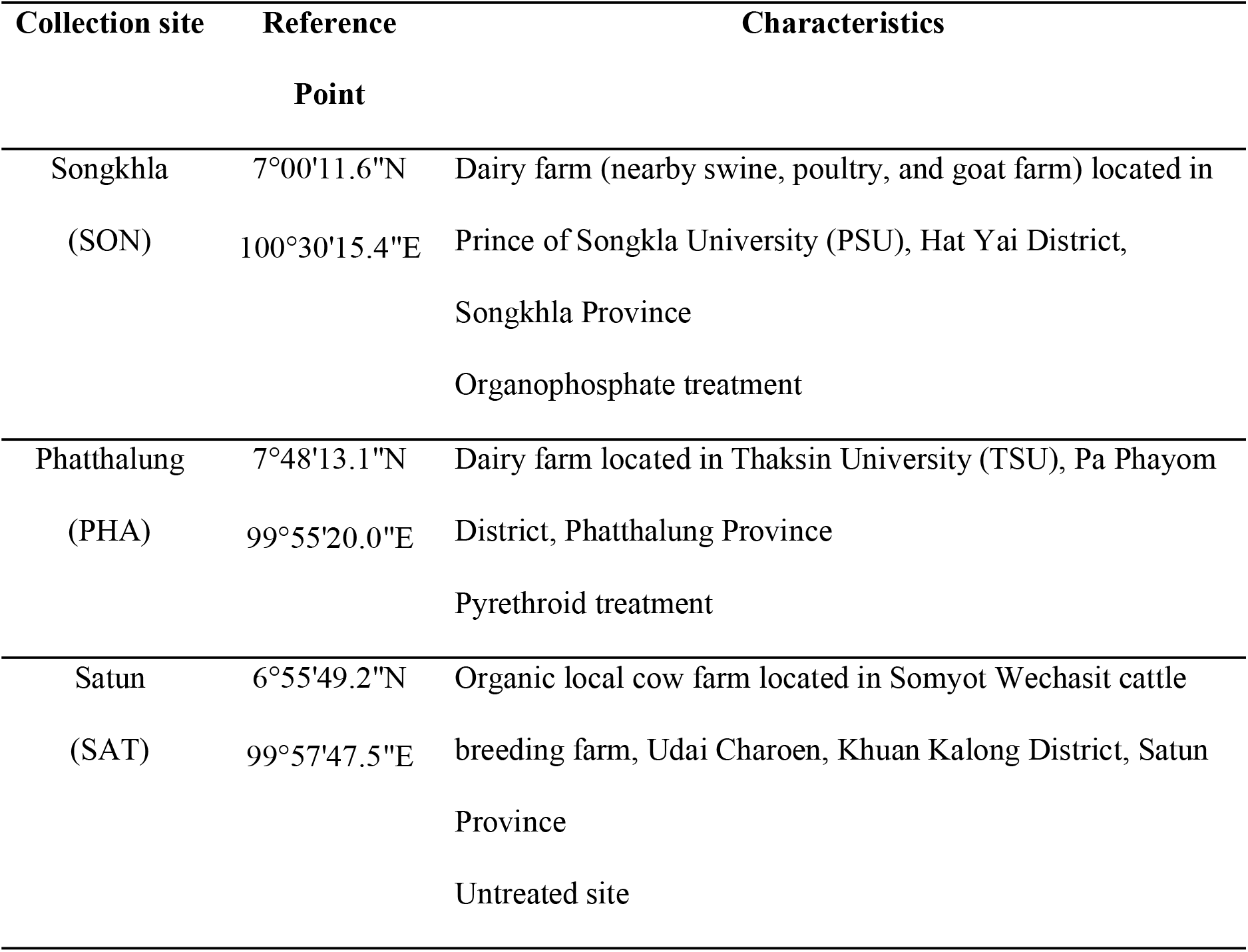
The collection sites of stable flies were located in three provinces: Songkhla (SON), Phatthalung (PHA), and Satun (SAT). The associated characteristics and insecticide use are summarized

### Insecticide-treated paper preparation

Insecticide-treated filter papers were prepared by using acetone solutions of synthetic pyrethroid insecticides, including a technical grade of permethrin (92.15% purity), deltamethrin (99.16% purity), alpha-cypermethrin (97.34% purity), cypermethrin (93.52% purity), lambda-cyhalothrin (97.16% purity), bifenthrin (98.35% purity) (Sherwood Chemicals Public Company Limited, Suan Luang, Bangkok) and commercial-grade cypermethrin (25.00% w/v) (Intergrade Trading Co. Ltd, Bangkapi, Bangkok). Absolute acetone (99.99%) was used as the solvent to dissolve each insecticide, and as the negative control. Rectangular sheets of filter paper measuring 10 × 10 cm (Whatman^®^ No. 1) were impregnated with 1 mL of each solution, based on the recommended concentration of each technical grade insecticide: permethrin (0.5500% w/v), deltamethrin (0.0500% w/v), alpha-cypermethrin (0.0750% w/v), cypermethrin (0.2500% w/v), lambda-cyhalothrin (0.0750% w/v) and bifenthrin (0.0625% w/v), and one commercial-grade insecticide cypermethrin (25.00% w/v) for residual treatment of resting-site for fly control (ODOC5 2018). The treated papers were stored at 4 °C and each was used only once.

### WHO cone bioassay for residual contact test

The tests were performed with slight modifications according to Tainchum et al., (2018). Insecticide susceptibility bioassays were performed using World Health Organization (WHO) cone test kits (WHO 2006). The WHO cone was placed on a filter paper for actual treatment or control. Five stable flies (both sexes) of each species were exposed to an insecticide-treated paper for 30 min and then transferred to clean plastic cups (5 individuals per cup), and provided with a cotton pad soaked with water and honey on the lid of the cup. Six replicates were carried out for each insecticide susceptibility test. The numbers of knocked-down (KD) flies were recorded, for a rapid knockdown at 30 min and a regular one at 60 min, and mortality was observed at 12 and 24 hr after treatment.

### Data analysis

The insecticide susceptibility status of stable flies was evaluated and interpreted following the revised WHO criteria (WHO 2016). The observed mortality was adjusted by Abbott’s formula when the control mortality was greater than 5% and less than 20% (Abbott 1925). The interpretation of percent mortality of tested stable flies at 24hr after insecticide exposure was labeled as one of susceptible (mortality 98-100%), incipient resistance (mortality 90-97%), and resistant (mortality below 90%) (WHO 2016). The responses of stable flies were compared between knockdown times (30 min and 60 min) and mortalities (12 hr and 24 hr) by calculating the differential knockdown time (DKT) = percent knockdown at 60 min - percent knockdown at 30 min and the differential mortality time (DMT) = percent mortality at 24hr - percent mortality at 12hr. A positive differential response indicates no recovery of the tested stable flies, while a negative value indicates recovery from knockdown or apparent mortality.

The mean values of percent knockdown and mortality were tested for normality of which the Shapiro-Wilk test (*P*>0.05) and were compared using an ANOVA at 5% of probability. The means were separated using Tukey’s significant difference test if the ANOVA statistic was significant (*P*<0.05). The analysis was performed using the SPSS software for Windows (version 10; SPSS Inc.; Chicago, IL, USA).

## Results

Pyrethroid susceptibility of two stable flies (*S. calcitrans* and *S. indicus*) was assessed by the WHO cone bioassay. This is an initial report on monitoring the distribution of pyrethroid insecticide effectiveness against wild stable fly populations in southern Thailand. The results of bioassays are shown in Tables 2–3. Overall, the knockdown at 60 min after exposure was in the range 73.33-100% across all tested populations of *S. calcitrans* and all insecticides, whereas *S. indicus* had 40-100% knockdown levels. The mortality range was from 26.67% to 95.83% and from 13.33% to 100% for *S. calcitrans* and *S. indicus*, respectively.

**Table 2.**
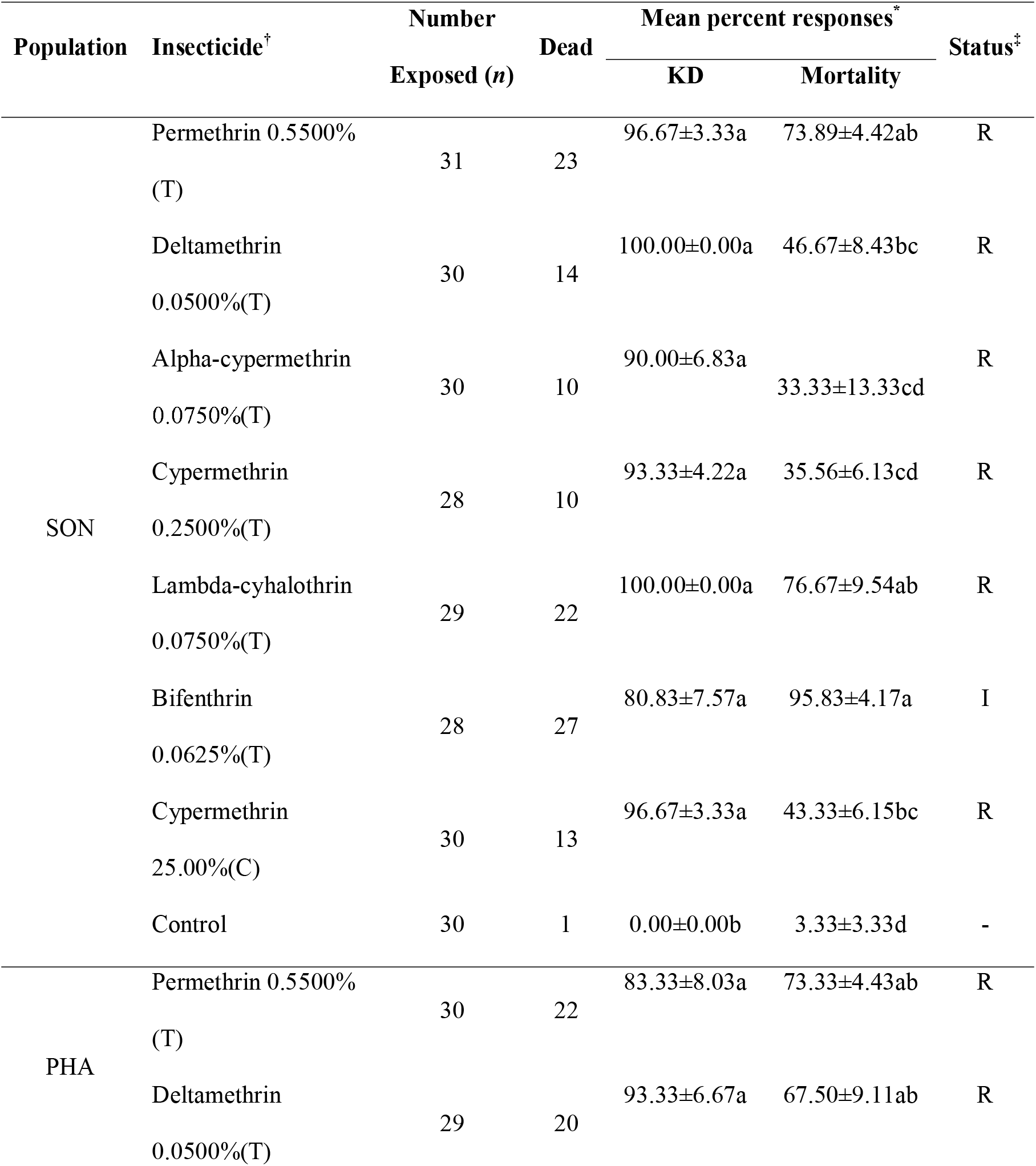
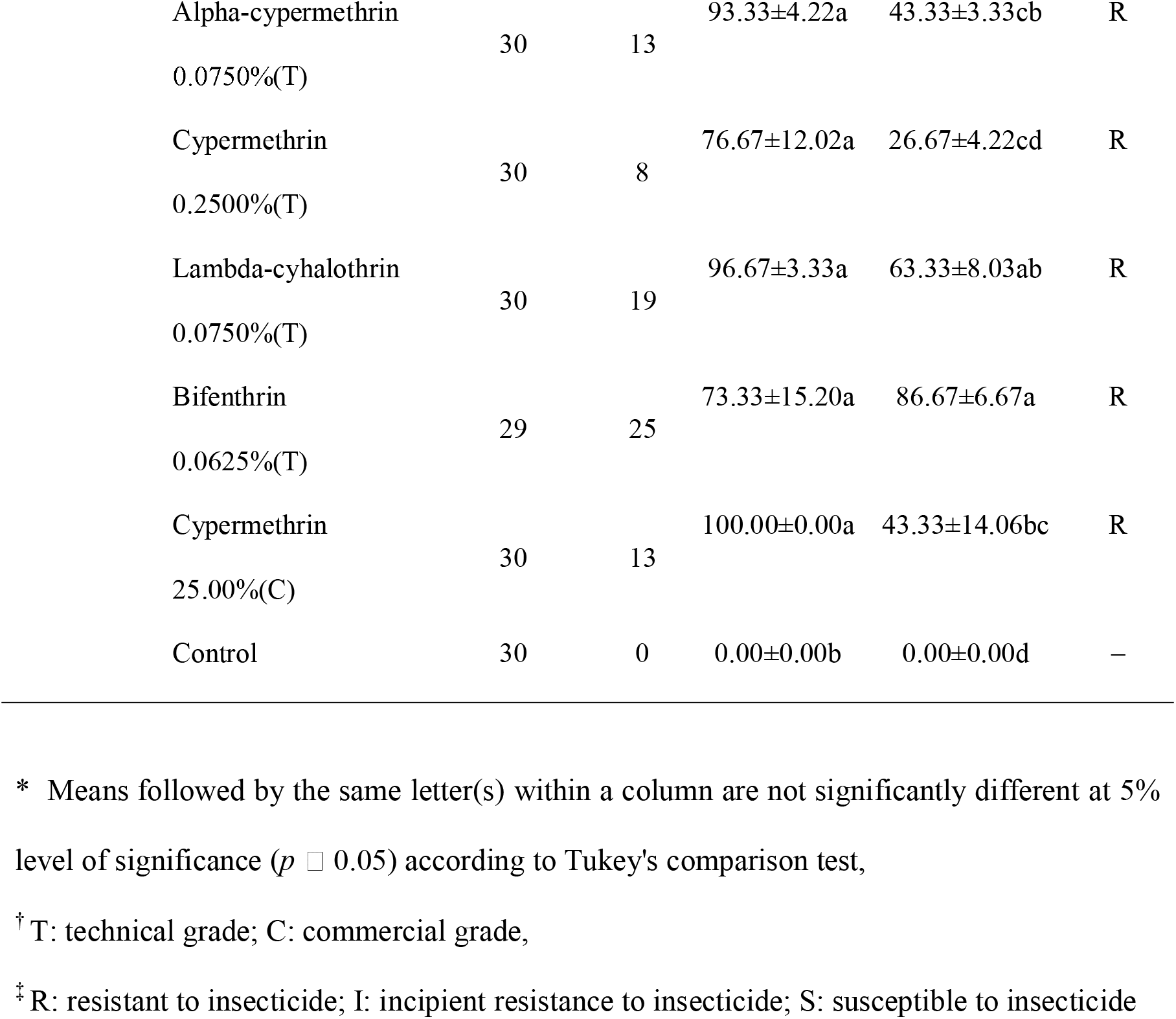
Wild adult *Stomoxys calcitrans* (males and females) sampled from Songkhla (SON) and Phattalung (PHA) populations, the mean knockdown (KD) and mortality rates (percentage ± SE) of within 60 min and 24h following 30 min exposure to recommended concentrations of six technical grade and one commercial-grade insecticide, in WHO cone bioassay

| Population | Insecticide<sup>†</sup> | Number<br>Exposed (n) | Dead | Mean percent responses<sup>*</sup> |  | Status<sup>‡</sup> |
| --- | --- | --- | --- | --- | --- | --- |
|  |  |  |  | KD | Mortality |  |
| SON | Permethrin 0.5500%<br>(T) | 31 | 23 | 96.67±3.33a | 73.89±4.42ab | R |
| | Deltamethrin<br>0.0500%(T) | 30 | 14 | 100.00±0.00a | 46.67±8.43bc | R |
| | Alpha-cypermethrin<br>0.0750%(T) | 30 | 10 | 90.00±6.83a | 33.33±13.33cd | R |
| | Cypermethrin<br>0.2500%(T) | 28 | 10 | 93.33±4.22a | 35.56±6.13cd | R |
| | Lambda-cyhalothrin<br>0.0750%(T) | 29 | 22 | 100.00±0.00a | 76.67±9.54ab | R |
| | Bifenthrin<br>0.0625%(T) | 28 | 27 | 80.83±7.57a | 95.83±4.17a | I |
| | Cypermethrin<br>25.00%(C) | 30 | 13 | 96.67±3.33a | 43.33±6.15bc | R |
| | Control | 30 | 1 | 0.00±0.00b | 3.33±3.33d | - |
| PHA | Permethrin 0.5500%<br>(T) | 30 | 22 | 83.33±8.03a | 73.33±4.43ab | R |
| | Deltamethrin<br>0.0500%(T) | 29 | 20 | 93.33±6.67a | 67.50±9.11ab | R |

|  |  |  |  |  |  |
| --- | --- | --- | --- | --- | --- |
| Alpha-cypermethrin | 30 | 13 | 93.33±4.22a | 43.33±3.33cb | R |
| 0.0750%(T) |  |  |  |  |  |
| Cypermethrin | 30 | 8 | 76.67±12.02a | 26.67±4.22cd | R |
| 0.2500%(T) |  |  |  |  |  |
| Lambda-cyhalothrin | 30 | 19 | 96.67±3.33a | 63.33±8.03ab | R |
| 0.0750%(T) |  |  |  |  |  |
| Bifenthrin | 29 | 25 | 73.33±15.20a | 86.67±6.67a | R |
| 0.0625%(T) |  |  |  |  |  |
| Cypermethrin | 30 | 13 | 100.00±0.00a | 43.33±14.06bc | R |
| 25.00%(C) |  |  |  |  |  |
| Control | 30 | 0 | 0.00±0.00b | 0.00±0.00d | – |

\* Means followed by the same letter(s) within a column are not significantly different at 5%
level of significance ( $p \leq 0.05$ ) according to Tukey's comparison test,
†T: technical grade; C: commercial grade,
‡R: resistant to insecticide; I: incipient resistance to insecticide; S: susceptible to insecticide

**Table 3.**
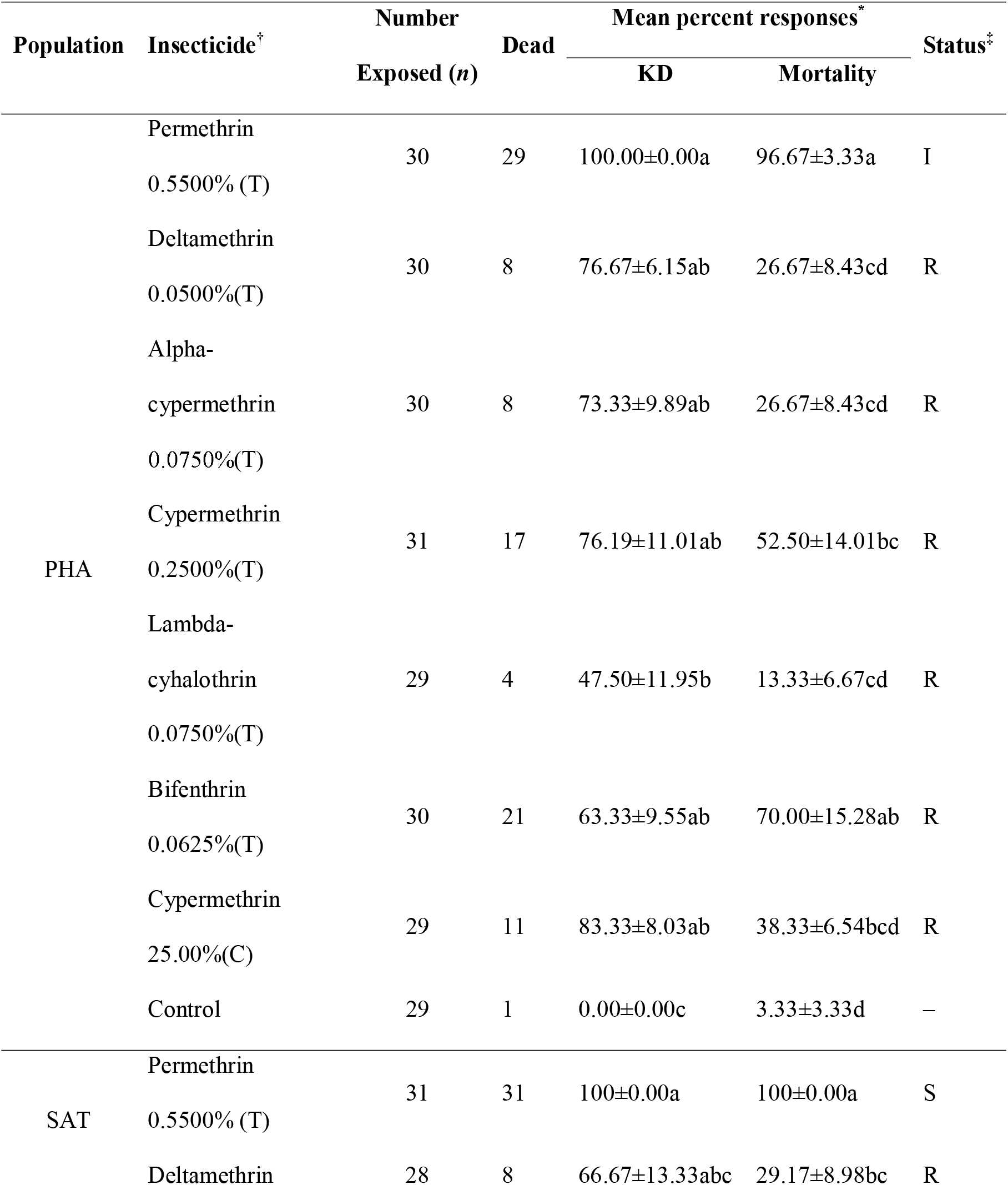
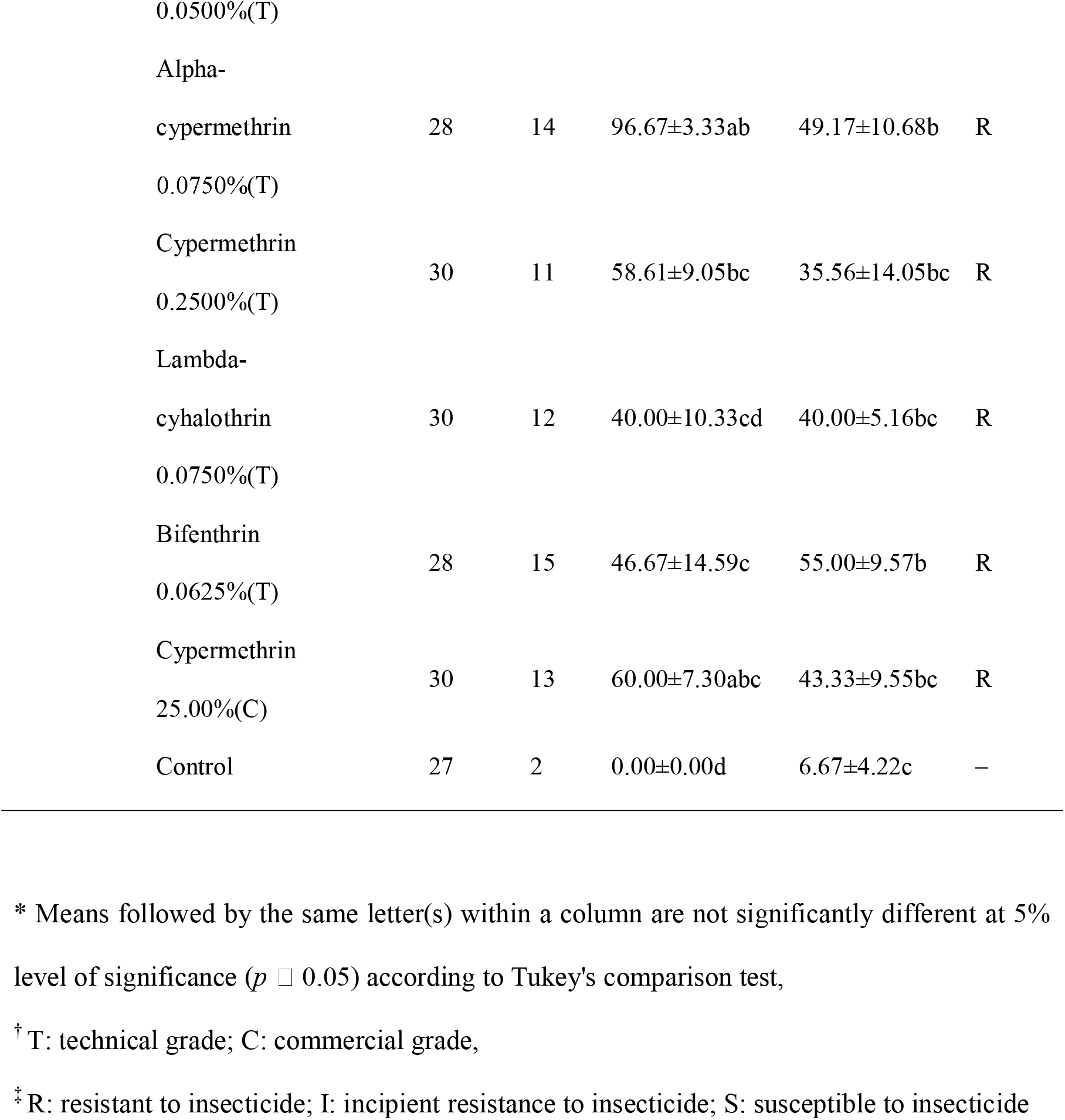
Wild *Stomoxys indicus* (males and females) adults from Phattalung (PHA) and Satun (SAT) populations, the percent knockdown (KD) at 60 min and 24h mortality (percentage ± SE) after exposure to an insecticide at recommended concentration, for six technical grade and a commercial-grade insecticide

The *S. calcitrans* SON population, exhibited a high level of knockdown by all the insecticides (>90%) except for bifenthrin that gave 80.83% knockdown. A complete knockdown was found in samples tested with deltamethrin and lambda-cyhalothrin. The highest mortality rate at 24 hr of tested *S. calcitrans* populations to a pyrethroid was found in the SON population with bifenthrin at 95.83%, while the mortality caused by other insecticides was below 90% (Table 2). Commercial cypermethrin provided complete knockdown (100%) of *S. calcitrans* in the PHA population. The high knockdown level (>90%) of *S. calcitrans* in the PHA population was found from exposure to deltamethrin, alpha-cypermethrin, and lambda-cyhalothrin. Similar to the SON strain, the highest mortality in samples of *S. calcitrans* was caused by bifenthrin (86.67% mortality) (Table 2).

For *S. indicus*, the knockdown observed at 60 min after exposure to insecticide was the highest in both PHA and SAT populations with permethrin. The SAT population when tested with alpha-cypermethrin had a high 96.67% knockdown rate, while relatively low knockdown levels were found with lambda-cyhalothrin in both PHA and SAT populations. The highest mortalities of both *S. indicus* populations were found when exposed to permethrin, namely 96.67% and 100% for samples collected from PHA and SAT, respectively. These were considered to have incipient resistance in the PHA strain while the SAT strain was resistant. There was no significant difference in mortality rates of PHA strain when tested with permethrin and bifenthrin, but they gave higher mortalities than the other insecticides (Table 3). Mortalities caused by all other insecticides in the SAT population, except for permethrin, were not significantly different. From these results, it is obvious that all tested stable flies (strains from all sampled locations) were resistant to all pyrethroid insecticides, except for *S. indicus* from both PHA and SAT when exposed to permethrin (Table 2–3).

The differential mortality responses of *S. calcitrans* by time (24hr and 12hr) was the largest in samples collected from PHA when exposed to bifenthrin (51.72%), while the differential in knockdown by time (60 min and 30 min) was the largest (82.14%) in samples collected from SON tested with bifenthrin (Fig. 2A–B). In *S. indicus*, the largest differential mortality found was in the PHA population exposed to bifenthrin (13.33%). Similar to *S. calcitrans*, the differential knockdown by time was also the highest in samples of *S. indicus* from PHA with bifenthrin (63.33%), followed by lambda-cyhalothrin (−34.48%) (Fig. 2C–D).

**Fig 2.**
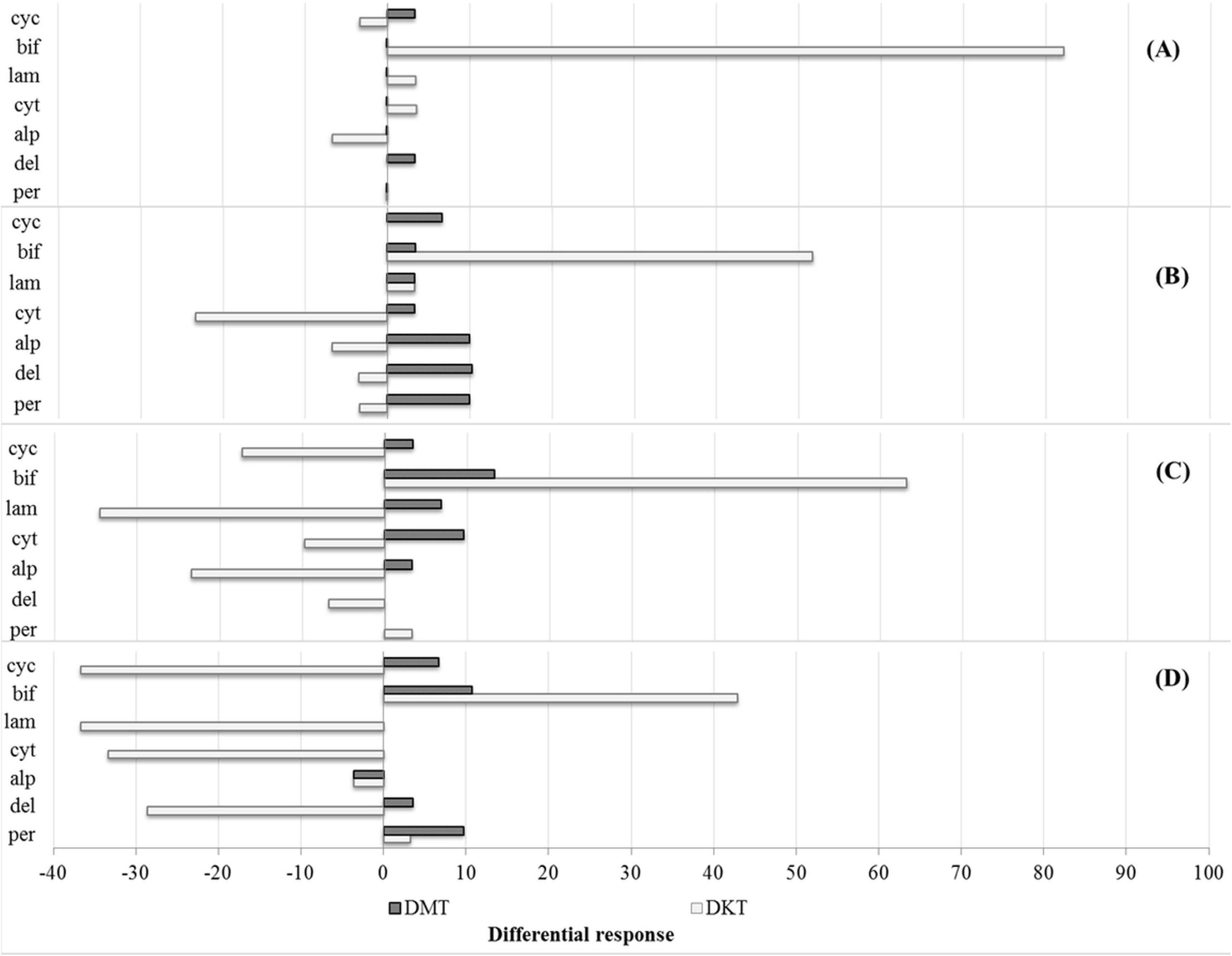
The differential response of *Stomoxya calcitrans* (A: Songkhla, B: Phatthalung) *Stomoxys indicus* (C: Phatthalung, D: Satun) exposed to pyrethroids (per=permethrin, del=deltamethrin, alp=alphacypermethrin, cyt=cypermethrin (tec.), lam=lambdacyhalothrin, bif=bifenthrin, cyc=cypermethrin (com.)) at two knockdown observations (60 min and 30 min after exposure) and mortalities (24hr and 12hr), DMT: differential mortality time, DKT: differential knockdown time (DMT)

### Comparability between the pyrethroid insecticide response of two species, four populations of stable fly

The pyrethroid insecticide response of four populations was performed by Turkey’s multiple range test (*P*<0.05)(Table 4). All four populations, devoid of species and province, were compared for the mortality rate (%). There was no significant difference in the mortality rate of stable flies exposed to alphacypermethrin, cypermethrin (Tec), cypermethrin (Com), and negative control. Completely susceptible to permethrin, 100% mortality was seen in *S. indicus* SAT which was significantly different with another 3 populations (73.33% mortality). While *S. indicus* SAT showed the significantly lowest mortality rates in the deltamethrin and bifenthrin were less than half of the *S.calcitrans* SON, *S.calcitrans* PHA and *S.indicus* PHA.

**Table 4.** Comparison of the mean number of pyrethroid insecticide response (mortality rates) of 4 populations of stable flies (*1=Stomoxys calcitrans* SON; 2= *Stomoxys calcitrans* PHA; 3= *Stomoxys indicus* PHA; 4= *Stomoxys indicus* SAT)

| Chemical | Test populations* |  |  |  | Statistic |
| --- | --- | --- | --- | --- | --- |
|  | 1 | 2 | 3 | 4 |  |
| Permethrin 0.5500% (T) | 73.33<sup>b</sup> | 73.33<sup>b</sup> | 73.33<sup>b</sup> | 100<sup>a</sup> | $F=2.196$<br>$P=0.120$ |
| Deltamethrin 0.0500%(T) | 46.67<sup>a</sup><sup>b</sup> | 67.50<sup>a</sup> | 67.50<sup>a</sup> | 29.17<sup>b</sup> | $F=3.278$<br>$P=0.042$ |
| Alpha-cypermethrin 0.0750%(T) | 33.33<sup>a</sup> | 43.33<sup>a</sup> | 43.33<sup>a</sup> | 49.17<sup>a</sup> | $F=0.317$<br>$P=0.813$ |
| Cypermethrin 0.2500%(T) | 35.55<sup>a</sup> | 26.67<sup>a</sup> | 26.67<sup>a</sup> | 35.55<sup>a</sup> | $F=2.069$<br>$P=0.137$ |
| Lambda-cyhalothrin 0.0750%(T) | 76.67<sup>a</sup> | 63.33<sup>ab</sup> | 63.33<sup>ab</sup> | 40.00<sup>b</sup> | $F=2.196$<br>$P=0.120$ |
| Bifenthrin 0.0625%(T) | 95.83<sup>a</sup> | 86.67<sup>a</sup> | 86.67<sup>a</sup> | 55.00<sup>b</sup> | $F=1.230$<br>$P=0.325$ |
| Cypermethrin 25.00%(C) | 43.33<sup>a</sup> | 43.33<sup>a</sup> | 43.33<sup>a</sup> | 43.33<sup>a</sup> | $F=23.621$<br>$P=0.000$ |
| Control | 3.33<sup>a</sup> | 0.00<sup>a</sup> | 0.00<sup>a</sup> | 6.67<sup>a</sup> | $F=0.000$<br>$P=0.000$ |

\* Means followed by the same letter(s) within a row are not significantly different at 5% level of significance ( $p \leq 0.05$ ) according to Tukey's comparison test.

## Discussion

Inappropriate and excessive use of insecticides causes contamination in the environment and induces resistance in arthropods. Their metabolites result in a reduction of contamination events and increase the challenge for those who are involved in livestock production, along with insecticide resistance issues (Brito et al. 2018). Adult stable flies can be exposed to insecticides that are used during outbreak situations, such as insecticidal fogging and residual spraying to reduce the impacts of stable flies on animals. Exposure can also occur through programs for controlling other livestock ectoparasites, such as house flies, *Musca domestica* Linnaeus (Olafson et al. 2019). The resistance of stable flies to commonly used insecticide classes has been previously reported for organochlorines, organophosphates, and pyrethroids in the U.S. and some European countries (Somme1958, Cilek and Greene 1994, Pitzer et al. 2010). Synthetic pyrethroids have been used extensively in Thailand for the control of agricultural and livestock pests, and also against disease vectors (Pakvilai et al. 2012, Chareonviriyaphap et al. 2013, Wongta et al., 2018). Resistance to pyrethroids was first reported in stable flies in Kansas cattle feedlots in 1994 (Cilek and Greene 1994) and the presence of resistant stable fly populations in another area was reported later on (Pitzer et al. 2010, Salem et al. 2012, Tainchum et al. 2018). No prior study of the susceptibility of stable flies to insecticides has been done in Thailand. No standard bioassay has yet been recommended for stable fly testing, unlike for the testing of mosquitos or house flies. In addition, this is the first report to study on insecticide susceptibility status of *S. indicus*. This fly species was found to be the second-most numerous species in all the study sites, and showed crepuscular activity in a previous report (Lorn et al. 2020); therefore, a study on susceptibility to insecticides of this species was worthwhile.

This study was carried out to investigate the status of pyrethroid susceptibility in stable flies sampled from different sites in the southern part of Thailand. Our findings show that *S. calcitrans* was the most susceptible to bifenthrin, while *S. indicus* was the most susceptible to permethrin. Bifenthrin is a non-alpha-cyano pyrethroid insecticide and an acaricide recommended by the WHO for indoor residual spraying. It has a relatively low irritant and knockdown effect on mosquitoes; thus allowing the mosquitoes to rest on treated surfaces for a longer period for exposure to a lethal dose, and hence provides a high mortality rate (Komalamisra et al. 2009). It is a newly synthesized pyrethroid and its effectiveness has been studied preliminarily in Thailand, in 1998 (OVBDC 1999). These properties could have an impact on the stable fly population density if complete spraying coverage is achieved in a community. The most permethrin susceptibility in this study was seen in *S. indicus*. Where a new combination of fipronil and permethrin provides excellent repellency and insecticidal efficacy for at least 5 weeks against *S. calcitrans* (Fankhauser et al. 2015). The knockdown resistance (*kdr*) associated with permethrin resistance was detected in stable flies from the United States, Costa Rica, France and Thailand (Olafson et al. 2019). Regarding the susceptibility to pyrethroids, the results from this current study demonstrate that the recommended concentration of any of the phenotypical tested pyrethroids (recommended concentration of commercial-grade) was not completely able to control the two field populations of *S. calcitrans*, while interestingly *S. indicus* in the Satun sample was still susceptible to permethrin. The collection site in Satun Province was located in an organic local cow farm, not treated with an insecticide. However, resistance to most tested pyrethroids was occurring in samples collected from this site. This may be explained by the dispersal of resistant stable flies that may happen more readily on nearby dairy farms or agricultural areas, where such selection pressure was present, impacting the effectiveness of chemical control at nearby dairies. The low susceptibility of stable flies to pyrethroids seen in this study suggests that current control practices relying on these insecticides are inadequate for further deployment. The recommended concentrations of pyrethroids for controlling stable flies should be adjusted to achieve proper control and elimination. However, the susceptibility of stable flies in other areas of Thailand has not been reported. The susceptibility of different populations of stable flies to insecticides may be affected by various factors, such as geographic variations, breeding habitat, health and age of stable flies, and pattern of insecticide use. Further investigations are needed to identify the susceptibility and the potential forces driving resistance in stable flies in Thailand.

The insecticide resistance of stable fly in this study may have consequences to the emerging of cattle and buffalo diseases along with southern Thailand, especially arthropod-borne viral (arboviruses) diseases -lumpy skin disease (LSD), meaning that the ordinary insecticide control program is not as effective as it could be. The first case occurred on 29 March 2021, an LSD outbreak involving beef cattle farms in Roi-Et Province, Northeastern Thailand (Arjkumpa et al. 2021). Then the outbreak was spread all around the country. The FAO team reported that the economic impact of LSD on South, East and Southeast countries was estimated to be up to USD 1.45 billion in direct losses of livestock and production.

Vaccination could assist to prevent this LSD but still an absence of large-scale vaccination targeting susceptible livestock (Roche et al. 2020). So the effective vector control can be a support to improve the reduction of disease transmission between host-vector contact.

Insecticide resistance develops with an evolutionary process of natural selection, which adapts that insect population to insecticides. The main mechanisms involve either mutation within the target site of the insecticide and/or an increased rate of insecticide detoxification (Nkya et al. 2013). To confirm insecticide resistance in a stable fly population, molecular techniques could assist in identifying the prevalence of such modifications in each population of stable flies. The recent report by Olafson et al. (2019) indicated that a low frequency of the *kdr* allele was observed in a stable fly population from the Wang Nam Khiao, Nakhon Ratchasima, Thailand. The resistance mechanisms should be elucidated to support estimating resistance and to design an effective strategy for the proper control of the target pests. Moreover, standard insecticide resistance detection assays for stable flies should be developed, with improved protocols, to facilitate collecting baseline information.

Further classes of insecticides should be examined and monitored for generating complementary data on the level of resistance, and possible evolution in the level of resistance in stable flies should be observed. The control of stable fly populations could be achieved by using an integrated approach or integrated vector management (IVM), with innovations in trapping, sanitization measures, parasitoid insects, botanical insecticides, and the application of efficient chemical compounds.

## Conclusion

The phenotypic pyrethroid insecticide susceptibility conclusively demonstrates mostly resistance in stable fly populations of the southern provinces in Thailand. This study indicates that the stable fly populations in Thailand should be further monitored, not only regarding the current insecticide used to protect livestock but also for new semiochemical toxicity and/or for different mechanisms of insecticide resistance, to improve the effectiveness of stable fly control. Also, new standard techniques for insecticide susceptibility detection in the stable fly samples should be considered, as these could be essential tools to consistently measure insecticide resistance.

## Acknowledgements

This research study was kindly supported by the National Science, Research and Innovation Fund (NSRF) and Prince of Songkla University (Grant No.NAT6505003c) and Program Management Unit (Area-based) (PMUA107470), Office of National Higher Education Science Research and Innovation Policy Council. We are grateful to the Research and Development Office, PSU, and Dr. Seppo Karrila for their helpful editing and proofreading services. The authors would like to express special gratitude to the cattle farms, which belong to the Faculty of Technology and Community Development, Thaksin University, Faculty of Agriculture, and to Mr. Somyot Wechasit, cattle breeding farm, Thailand, for their contribution to the fly collection.

